# Neural Tracking of Linguistic Predictors in Spontaneous Conversational Speech

**DOI:** 10.64898/2026.05.13.722865

**Authors:** Maïwenn Fleig, Shuai Wang, Anna E. Dudek, Jean-Marc Freyermuth, Leonor Becerra, Philippe Blache

## Abstract

This study investigates whether neural tracking of linguistic information extends from read speech to spontaneous conversation. Using the temporal response function (TRF) framework, we validate our approach on a read-speech EEG dataset and then apply it to EEG recordings from natural conversations. We observe reliable neural tracking of key linguistic predictors, including word onset, part-of-speech surprisal, and lexical surprisal, in spontaneous speech, with effects around 200, 400, and 600 ms. These results provide new evidence that linguistic neural tracking operates in natural conversational settings and confirm the feasibility of EEG studies in ecologically valid contexts.

## Introduction

The study of the neural bases of language comprehension relies on a range of techniques aimed at identifying the impact of specific linguistic phenomena in the brain signal. A substantial body of work has investigated different levels of linguistic processing by linking them to event-related potentials (ERPs) emerging at distinct time windows in the brain signal after word onset. More recently, a complementary approach has been proposed, known as the temporal response function (TRF), which consists in predicting the brain signal from linguistic predictors (Brodbeck et al., 2023; Crosse et al., 2016; Ding & Simon, 2012).

The temporal response function framework enables the fine-grained investigation of linguistic predictors, individually or jointly, by modeling the correspondence between continuous linguistic signals and brain activity over extended time scales. Using this approach, previous studies have demonstrated neural tracking of the speech envelope (Brodbeck & Simon, 2020; Ding et al., 2014) as well as of multiple linguistic predictors, including phoneme and word onsets, lexical surprisal, and semantic dissimilarity, highlighting the flexibility of TRFs for probing different levels of language processing (Broderick et al., 2018; Chalehchaleh et al., 2025; Gillis et al., 2021; Heilbron et al., 2022; Weissbart et al., 2020).

However, most TRF studies rely on passive listening to read speech under controlled EEG conditions, typically using audiobook stimuli. In contrast, only a few studies have examined natural conversational speech (Goldstein et al., 2025; Silem et al., 2025; Zada et al., 2024), largely due to the challenges of collecting and analyzing EEG data in ecological settings where speech production and movement introduce substantial noise.

In this study, we address the challenge of analyzing the neural correlates of speech in natural settings by investigating whether findings from read speech generalize to spontaneous speech. In the naturalistic setting, the brain not only processes incoming speech but also plans upcoming responses, engaging additional neural systems. It is also harder to predict due to disfluencies and greater variability in pacing/repairs. We hypothesize that principal linguistic predictors contributing to neural tracking are also active during spontaneous speech.

First, we introduce a processing pipeline that trains models on single linguistic predictors and validate it on an existing read-speech dataset (Bhattasali et al., 2020), successfully replicating previously reported results. Second, we apply the same pipeline to a corpus of spontaneous conversational speech (Boudin et al., 2023).

Our results confirm neural tracking of several linguistic predictors, including word onset, part-of-speech surprisal, and lexical surprisal in spontaneous speech with robust effects in canonical linguistic time windows around 200, 400, and 600 ms. To our knowledge, this study provides the first evidence of linguistic neural tracking in spontaneous speech and demonstrates the feasibility of using EEG data collected in naturalistic conversational settings

## Related works

One major advantage of TRFs is their ability to capture brain activity over extended time periods, representing a substantial methodological advance for the analysis of natural language. However, TRF studies have so far relied almost exclusively on read speech (Broderick et al., 2018; Chalehchaleh et al., 2025; Dou et al., 2025; Gillis et al., 2021; Heilbron et al., 2022; Weissbart et al., 2020). In the present study, we seek to advance our understanding of the neural bases of language in ecological contexts by focusing on spontaneous rather than read speech.

Predictor selection is a central issue in TRF studies. Extensive prior work has highlighted the crucial role of speech envelope tracking in neural responses (Brodbeck & Simon, 2020; Ding et al., 2014; Lalor & Foxe, 2010), demonstrating effects of rhythmic structure and acoustic onsets (Ding & Simon, 2014), as well as higher-level predictors such as phoneme onsets (Brodbeck et al., 2023; Donhauser & Baillet, 2020). These predictors typically elicit a negative deflection around 100 ms, sometimes followed by a second negativity around 250 ms, resembling the N250 or phonological mismatch negativity (Dou et al., 2025; Gillis et al., 2021).

At higher linguistic levels, word-based predictors such as word onset, surprisal, and semantic dissimilarity evoke later and distinct neural responses depending on their representational level. For instance, part-of-speech (POS) surprisal has been linked to effects in the 200 – 500 ms time window (Heilbron et al., 2022). At the lexical level, predictors including word onset, lexical surprisal, and word frequency are typically associated with responses around 400 ms, often interpreted as reflecting an N400 component (Broderick et al., 2018; Dou et al., 2025; Weissbart et al., 2020). Semantic-level predictors have also been investigated, most notably semantic dissimilarity (Broderick et al., 2018). These studies likewise report N400-like effects, although such findings are not always consistently replicated (Gillis et al., 2021).

Recent work has examined brain responses in terms of both latency and duration, highlighting the distinct temporal profiles of individual predictors — not only in their predictive power (Dou et al., 2025), but also in their position within the hierarchy of linguistic processing (Gwilliams et al., 2025).

## Methods

### Data

We used two datasets, to contrast a more controlled, passive listening scenario with a richer, more naturalistic and dynamically interactive conversational setting.

The first corpus is the Alice dataset (Bhattasali et al., 2020; J. R. Brennan, 2023), which includes EEG recordings from 49 participants listening to the opening chapter of Alice in Wonderland (12.4 min; segmented into 12 trials). The data was recorded using 61 electrodes at 500 Hz. Several participants had already been excluded by the original authors due to experimental errors, unmet behavioral criteria, or excessive noise, leaving 33 participants for the analysis. We excluded four additional participants because their first trial was missing.

For the second dataset, we used the SMYLE corpus (Boudin et al., 2023), a French multimodal dataset combining audio, video, and EEG recordings. It includes 30 dyads (16h) who first engage in storytelling and then in free conversation. Storytelling consists of three tasks: recounting a short video (the Pear Story (Chafe, 1981)), pitching a movie, book, or video game, and describing a memorable vacation. Two listener conditions were employed: attentive, in which listeners followed and responded naturally, and distracted, in which listeners secretly counted words starting with /*t*/. For this study, we selected 19 dyads after excluding dyads with excessive noise and focused on the storytelling task, in which one participant acts solely as listener to reduce EEG noise. The corpus provides enriched orthographic transcriptions (Blache et al., 2017), segmented into Inter-Pausal Units (IPUs) and annotated for laughter, disfluencies, repetitions, truncated words, and elisions. Transcriptions were normalized, tokenized, and time-aligned with the speech signal using the SPPAS toolkit (Bigi, 2012). EEG data were recorded with two 64-channel BioSemi systems (10 – 20 layout) at 2048 Hz.

### EEG Pre-processing

EEG preprocessing was conducted using MNE-Python 1.10 (Gramfort, 2013). Alice dataset bad channels were pre-marked by the authors (Bhattasali et al., 2020; J. R. Brennan, 2023). For SMYLE participants, noisy or artifact-ridden channels were marked as bad via visual inspection of raw signals and power spectra. Participants with >20% bad channels were excluded, resulting in two Alice and four SMYLE participants removed. Signals were referenced to the common average, band-pass filtered (0.5 – 30 Hz, FIR), and bad channels interpolated using spherical splines. To limit interpolation to <15% (Crosse et al., 2021), three electrodes were excluded per dataset, leaving 58 for Alice and 61 for SMYLE. EEG signals were downsampled to 256 Hz. We investigated broad band frequencies as well as frequencies from the delta band. We included the delta band, since word-related speech features naturally occur at 1–4 Hz, matching the temporal dynamics of delta oscillations

For the SMYLE dataset, recorded during natural conversations, a semi-automatic artifact removal procedure using ICA was applied. Signals were scaled to unit variance and whitened via PCA, then FastICA (Hyvarinen, 1999) extracted ICs according to the data rank. The ICLabel method was used to inspect and classify ICs as eye-blink, muscle, or cardiac artifacts (Li et al., 2022; Pion-Tonachini et al., 2019). In parallel, a human inspector classified ICs by visually examining IC time-series, topography and power spectrum. After excluding the ICLabel- and human-identified noise ICs, the EEG signals were reconstructed using all of the remaining ICs.

### Linguistic Features

We aim to investigate the neural correlates of a set of linguistic features, including higher-level features such as word and part-of-speech (POS) surprisal as well as low-level features like word onset and the speech envelope.

#### Surprisal Estimation using LLMs

The word surprisal and the POS surprisal were estimated using LLMs: GPT-2 (Radford et al., 2019) for Alice and GPT-fr (Simoulin & Crabbé, 2021) fine-tuned on French conversation for SMYLE. We argue that English LLMs are better equipped to model spoken English due to the substantially larger volume of available data and greater representation of spoken language compared with French. To address this gap, we fine-tuned the GPT-fr base model on a SMYLE-derived conversational dataset with LoRA applied to all layers, using transcriptions from both the storytelling and free-conversation tasks with all disfluencies preserved. The model was trained on samples of 10 consecutive turns separated by the `<p>` marker. Training ran for five epochs with AdamW, using the following parameters: learning rate = 0.002, 500-step warmup, batch size = 8, LoRA rank = 32, *α* = 32, dropout ratio = 0.05, and gradient clipping at 1.

We passed the transcriptions (of the first Alice chapter or the concatenated turns of the speaker for SMYLE) through the LLM and obtained the logits for each word. These were transformed into conditional probabilities for each word by applying softmax and choosing the most probable word. We obtain the word surprisal as follows:

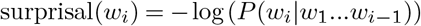

For the POS surprisal, we followed to approach of Heilbron et al. (2022), i.e. top-k nucleus sampling with *k* = 40 and *p* = 0.9. Specifically, the candidate set consisted of the most probable tokens whose cumulative probability reached 90%, with a minimum of the top 40 tokens always included. We restricted the maximum *k* to be 300 to reduce computational costs. We then calculated the POS tag using the Spacy library^1^ for each of the top-k nucleus sampled tokens as well as the actual target word. The POS surprisal is given by:

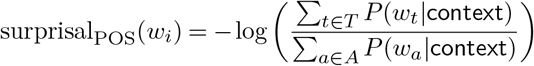

with *T* as the group of top-k nucleus sampled words having the same POS tag as the target word *w*_*i*_ and *A* as the group of **all** top-k nucleus sampled word. All words were derived given the same `context` sequence as *w*_*i*_.

#### Constructing Continuous Feature Signals

With our approach described above, we get discrete scores at each word onset. Since TRFs work on signals, we need to construct a continuous signal from these discrete values. For this, we initiate a continuous time series, or rather an array with the sampling rate of 256 Hz matching the duration of the conversation, set to zero throughout. Spikes scaled by the previously estimated surprisal value of the corresponding word were inserted at word onset times provided by the corpora. This yields a continuous surprisal representation. This procedure was repeated for the POS surprisal.

Because surprisal impulses occur at word boundaries, we additionally modeled a word onset feature to dissociate neural responses to linguistic information from responses driven purely by boundary timing. This control regressor was generated using the same procedure, but with impulses of amplitude 1 placed at each onset.

For the envelope, the amplitude envelope was extracted using the Hilbert transform implemented in the Eelbrain toolbox (Brodbeck et al., 2023), and then resampled to the target sampling rate of 256 Hz.

No normalization was applied to word onset since this feature was binary encoded. As suggested in (Crosse et al., 2021), the envelope was normalized by its standard deviation to maintain positive values. Given that surprisal and POS surprisal have identical timings as word onset, the non-zero values of the two features were z-score normalized.

### TRF Modeling

Temporal response functions are widely used to study the relationship between linguistic predictors and EEG signal by modeling how an input predictor, when convolved with a response function, predicts brain activity.

In practice, models are trained separately for each participant/dyad and feature, estimating time-lagged parameters that capture how predictors contribute to the EEG signal at different latencies. Both, feature and neural signal were normalized before training (except for word onset). Model performance is then evaluated by comparing the predicted EEG signal with the observed data across electrodes.

The estimated parameters, or weights, indicate how variations in the stimulus relate to EEG signals, reflecting the strength of coupling between the linguistic input and the neural response. When this coupling is strong, it is informative to examine whether its temporal profile corresponds to known ERP components. For instance, a strong coupling between lexical surprisal and EEG activity around 400 ms after stimulus onset may be interpreted as reflecting an N400-like effect.

In our study, the temporal lag range was defined from -200 ms to 800 ms, yielding 256 discrete time points given a sampling rate of 256 Hz. TRF weights were computed by minimizing the mean squared error between the recorded EEG signal and its model-based prediction. To reduce the risk of overfitting, for each dyad/participant, the EEG signals and the corresponding features were split into training and testing sets. For each participant in the Alice dataset, the first 10 trials (83% of the data) were used for training, the last two trials were reserved for testing. For each dyad in the SMYLE corpus, the initial 90% of the data was used for training and the remaining 10% was reserved for testing. TRF models were fitted separately on the training set of each dyad/participant using ridge regression, with parameter *λ* being optimized via 5-fold cross-validation, during which 100 candidate values logarithmically spaced between 10^−4^ and 10^12^ were evaluated. Model performance was quantified on the test set using the Pearson correlation coefficient of the predicted and observed EEG signals.

## Results

For both datasets, we trained TRF models for each participant using individual linguistic predictors. Overall model performance is summarized in Table 1. TRF significance was assessed with a mass univariate spatiotemporal cluster-based permutation test (Maris & Oostenveld, 2007) using MNE-Python’s `spatio_temporal_cluster_1samp_test`. The method computes univariate *t*-statistics across electrodes and time points, clusters adjacent points exceeding *p* = 0.01, and compares each cluster’s mass (sum of *t*-values) against a permutation-based null distribution of maximum cluster mass (10, 000 iterations) to determine significance (*p* < 0.05).

**Table 1.** Overall prediction accuracy (mean Pearson’s r) of TRF models for each feature for either broad (0.5 −30 Hz) or delta (0.5 − 4 Hz) bands.

| EEG Band | Feature | Pearson’s $r$ (Alice) | Pearson’s $r$ (SMYLE) |
| --- | --- | --- | --- |
| Broad | Envelope | 0.0395 | 0.0198 |
|  | Word Onset | 0.0486 | 0.0164 |
|  | POS Surprisal | 0.0131 | -0.0032 |
|  | Surprisal | 0.0017 | -0.0018 |
| Delta | Envelope | 0.0539 | 0.0252 |
|  | Word Onset | 0.0663 | 0.0235 |
|  | POS Surprisal | 0.0223 | -0.0050 |
|  | Surprisal | 0.0072 | 0.0011 |

### Envelope

Many studies have described acoustic neural tracking, in particular showing the predictive power of the speech envelope (Crosse et al., 2016; Ding & Simon, 2014; Drennan & Lalor, 2019; Yasmin et al., 2023). These works report several early effects starting as early as 50 ms and extending up to 250 ms. These effects are often associated with the N1–P2 complex, known to encode speech information. With regard to neural tracking, different studies have shown amplitude peaks at 50 – 80 ms (Ding et al., 2014) and at 130 – 200 ms (Drennan & Lalor, 2019; Yasmin et al., 2023). Our results (see Figure 1) on the Alice corpus show a similar double peak: the first one very early, around 30 ms (*p* < 0.019), and the second at about 170 ms (78 - 164ms, *p* < 0.048). This double peak therefore confirms a well-known effect in the neural tracking of read speech.

**Figure 1.**
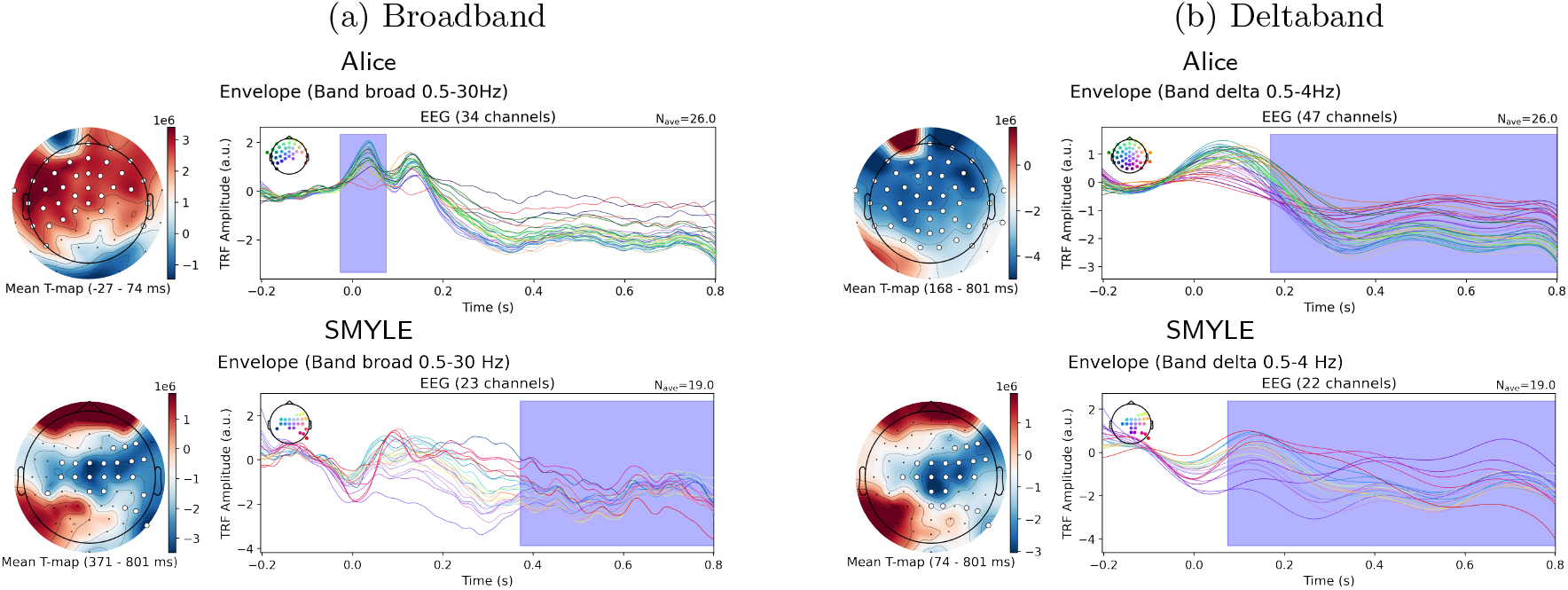
Spatiotemporal clusters were identified using a permutation-based approach for the **envelope**. *Left*: Scalp topography of the T-map averaged across significant post-stimulus time window. Electrodes belonging to significant clusters are marked by white circles, with the color scale indicating the magnitude of the statistics. *Right*: Averaged time-course of the envelope responses. The blue shaded region indicates the significant time window corresponding to the topographic map.

Our results in natural conversation show a pattern that could correspond to an amplitude modulation in the same temporal window, between 30 and 200 ms, showing the N1–P2 complex. A significant cluster only appears around 400 to 700 ms.

By comparing the results obtained for acoustic tracking in the delta band (0.5 - 4 Hz), which is frequently used in this field (Weissbart et al., 2020), we instead observe a greater similarity between the conditions (Figure 1b; *p* < 2 ×10^−4^ for Alice; *p* < 3 ×10^−3^ for SMYLE). In both cases, we indeed observe an overall comparable pattern across the entire time window studied, with a marked amplitude between 30 and 200 ms, this amplitude being slightly earlier and sharper in the case of read speech compared to spontaneous speech.

### Word Onset

Word onset is an important predictor for neural tracking. The literature reports an amplitude increase from 50 to 300 ms after word onset, and a second peak around 300 to 400 ms, usually interpreted as an N400 (Gillis et al., 2021; Karunathilake et al., 2023). In our results (see Figure 2), for read speech we observe a double-peak phenomenon around 150 ms and 230 ms, which then gradually decreases up to 500 ms (*p* < 0.0014). This observation doesn’t exactly match the findings in the literature regarding the second peak, which appears earlier here. This early effect, more pronounced around 200 ms, can possibly be associated with attentional phenomena typically observed in this time window (usually the P200). Results for spontaneous speech, on the other hand, show a peak around 300 – 500 ms (*p* < 0.05). This second observation corresponds more clearly to an interpretation in terms of the N400. In both cases, even though this phenomenon is less marked for read speech, it is interesting to note the strong predictive power of word onset in the 300 – 500 ms time window in both conditions (even in the absence of a clear peak for read speech). By examining the neural tracking of the word onset in the delta band, the results show a clearer peak between 150 and 350 ms for read speech (*p* < 7 × 10^−4^) and between 150 and 450 ms for spontaneous speech (*p* < 0.002). In this case, we observe more similar patterns across both conditions over a relatively broad shared time interval. In both cases, a peak around 300 ms is observed, which has been reported in studies describing a P300–N400 complex related to categorization and surprise.

**Figure 2.**
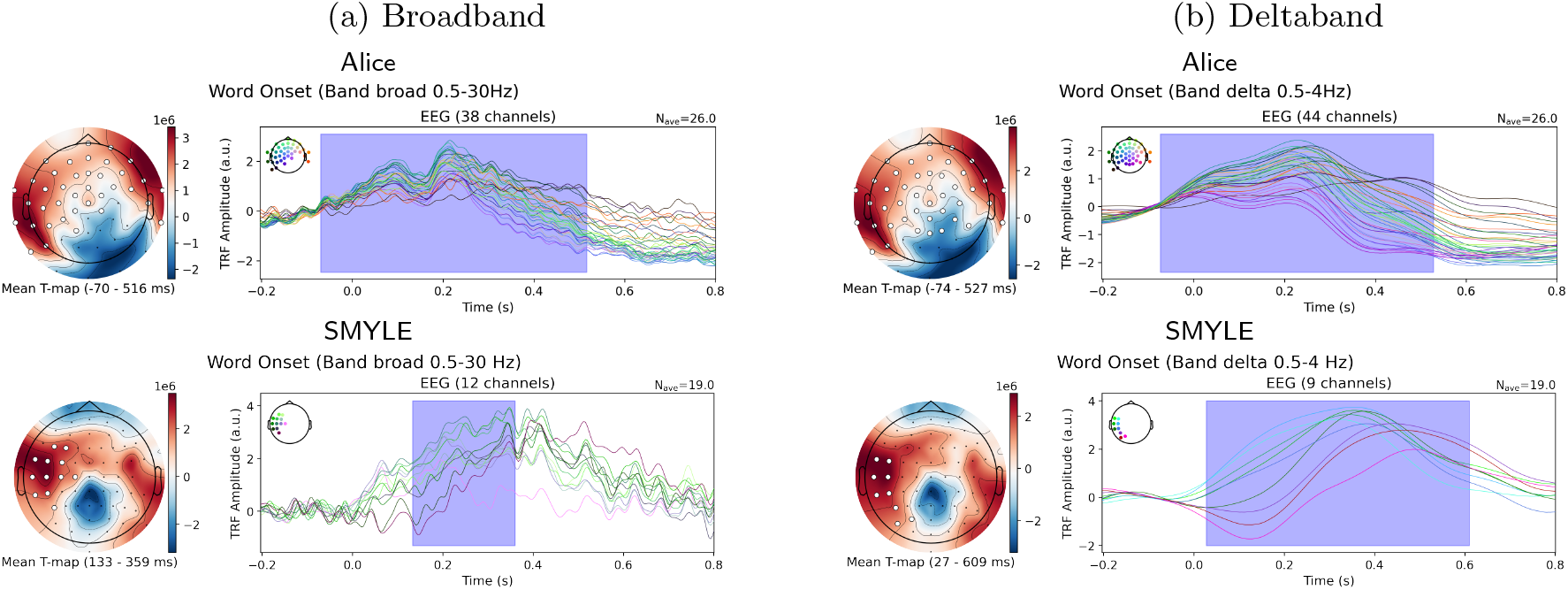
Spatiotemporal clusters of the weights trained on **word onset** responses were identified using a permutation-based approach. Significant clusters are highlighted by the topography and the blue shadow in the time courses.

### POS Surprisal

POS surprisal as a neural predictor is expected to show effects between 200 and 500 ms (J. Brennan & Hale, 2019; Heilbron et al., 2022). The effects associated with this predictor are more specifically syntactic, with a prominent impact around 400 ms, corresponding to effects observed in the P300–N400 complex related to prediction and memory access (Bornkessel-Schlesewsky & Schlesewsky, 2019). In addition, a later effect around 600 – 800 ms has also been reported (Heilbron et al., 2022). We observe similar effects starting around 100 ms (82 - 129 ms, *p* < 0.045), with a significant peak at 400 ms (see Figure 3; *p* < 0.02) and extending into a later effect in the Alice dataset (473 - 800ms; *p* < 9 × 10^−4^). Interestingly, the same pattern is also observed for spontaneous speech. Focusing more specifically on the delta band reveals the same POS surprisal impact pattern within the 200 – 700 ms time window (*p* < 0.0025 for Alice), with a comparable pattern observed for both read and conversational speech.

**Figure 3.**
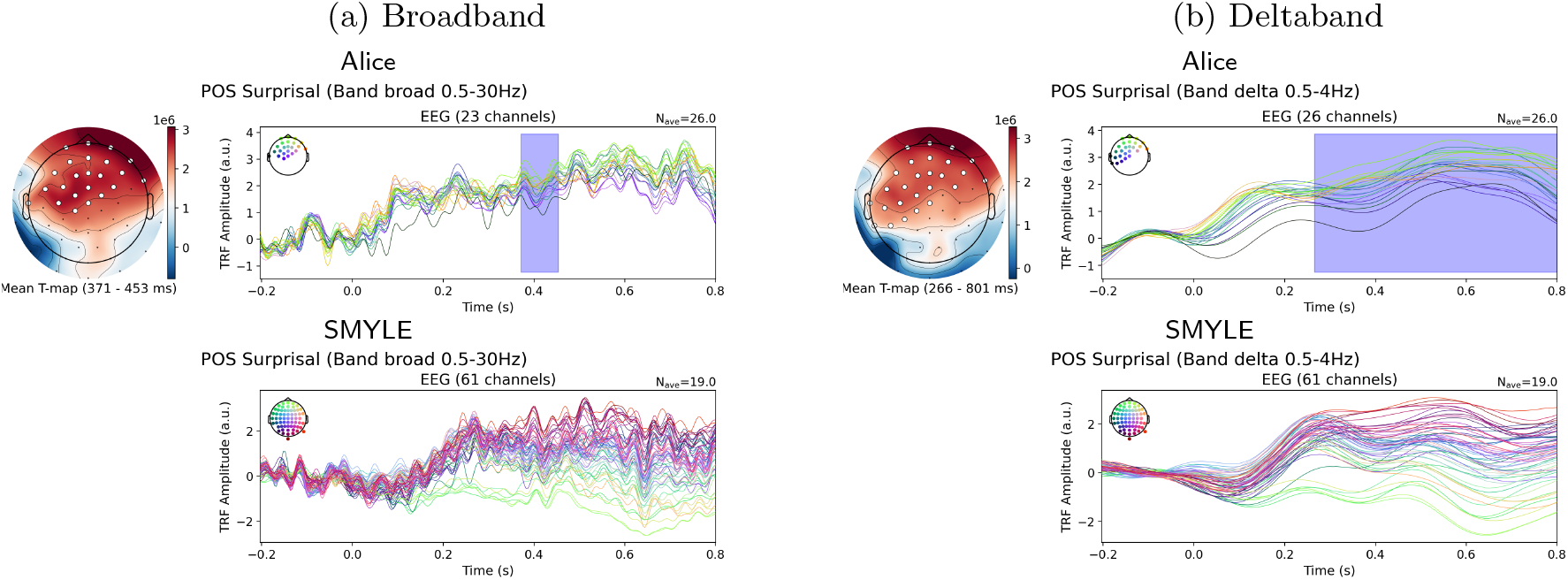
Spatiotemporal clusters of the weights trained on **POS surprisal** responses were identified using a permutation-based approach. Significant clusters are highlighted by the topography and the blue shadow in the time courses.

### Surprisal

The question of neural tracking of surprise has been investigated in several studies using read speech. These studies have generally shown an effect of lexical surprisal around 400 ms (Chalehchaleh et al., 2025; Gauthier & Levy, 2023; Gillis et al., 2021; Heilbron et al., 2022; Weissbart et al., 2020). This is an expected effect, which can be associated with the N400 component in ERP terms. It should be noted, however, that in these studies the results were characterized by small effect sizes and were often limited to a small number of electrodes. In addition, one study showed that multiple frequency bands can be shaped by surprisal, with in particular late effects observed between 200 and 1000 ms in the beta and gamma frequency bands (Weissbart et al., 2020).

Our results are more contrasted (see Figure 4). For the Alice dataset, as in the literature, we observe a slight impact of surprisal between 250 and 450 ms, although this effect is not very pronounced, thus confirming the limited effects reported in previous work. This effect appears in a more diffuse manner for spontaneous speech, with a possible impact starting around 350 ms. However, our results also show a significant late effect around 700 ms (*p* < 0.02). Interestingly, these findings are very different from those obtained with the word onset predictor, which may seem surprising. The analysis of neural tracking in the delta band does not allow us to draw comparable and significant conclusions between read and spontaneous speech either. Nevertheless, a pattern can be observed for read speech, with a first effect around 150 ms and a second around 300 ms, which is not visible for spontaneous speech.

**Figure 4.**
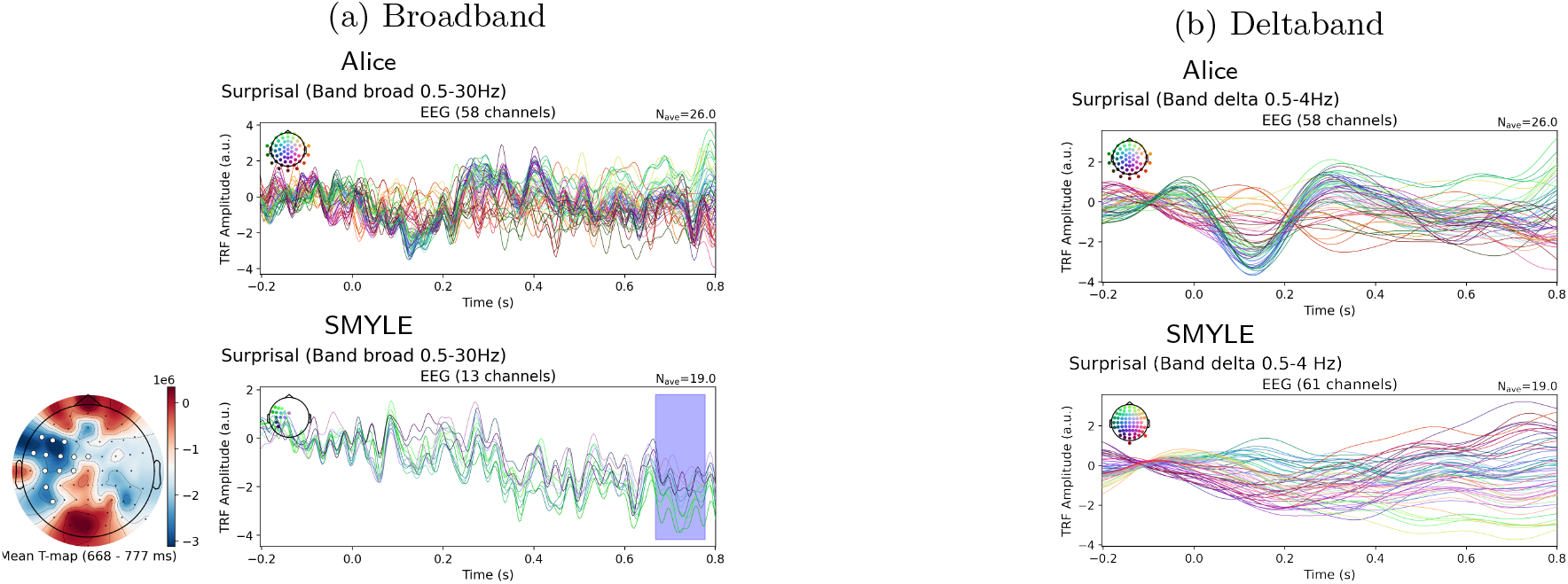
Spatiotemporal clusters of the weights trained on **surprisal** responses were identified using a permutation-based approach. Significant clusters are highlighted by the topography and the blue shadow in the time courses.

## Discussion & Conclusion

As expected, TRF models trained on read speech show higher predictive performance than those trained on naturalistic spontaneous speech (Table 1). In both contexts, linguistic features exhibit a similar performance pattern: low-level features outperform high-level features, and restricting neural tracking to the delta band substantially improves model performance. Unlike the narrative speech in Alice (audio), natural conversation in SMYLE provides intensive multimodal input, requiring the brain to integrate visual, auditory, linguistic, and social information. This environmental complexity may increase EEG variability, including greater spatial variation across electrodes and temporal asynchrony, resulting in fewer significant clusters in SMYLE than in Alice. However, when analysis is restricted to the delta band, typically associated with word-level speech frequency, the results show remarkable consistency between Alice and SMYLE (see Figure 1 and 2).

Our objective was to assess the feasibility of studying neural tracking in spontaneous speech. We used temporal response functions, which rely on extracting linguistic predictors from speech signals. Here, we employed commonly used predictors spanning low-level features (speech envelope, word onset) and higher-level features (part-of-speech and lexical surprisal). To this end, we fine-tuned a large language model on spontaneous speech to extract predictors tailored to this setting and used them to predict EEG signals.

Our results first allowed us to replicate findings from the literature for these predictors using a read-speech dataset. Importantly, results from spontaneous speech revealed comparable neural tracking effects. Together, these findings pave the way for new investigations of spontaneous speech, complementing the controlled speech paradigms traditionally used in neurolinguistics.

Further studies using multiple frequency bands are expected to discriminate different cognitive processes in natural conversations.Additionally, it would be valuable to examine predictors using multivariate models to assess the unique variance contributed by each feature. Future research should also explore the use of models trained on larger corpora or multilingual models to assess whether these factors can improve performance and generalisability.

## Footnotes

1 https://spacy.io/ with `fr_core_news_lg` for French and `en_core_web_lg` for English.

